# Which author is which? Gender Authorship Position in Aquaculture Literature

**DOI:** 10.1101/486209

**Authors:** Morgan Chow, Hillary Egna, Jevin West

## Abstract

Examining authorship position in aquaculture facilitates an improved understanding of status of women in the discipline, as authorship is a critical factor in professional success. In a review of more than eight million papers in the JSTOR Corpus across disciplines, West et al. 2013 found that men predominate in the first and last author positions and women are underrepresented in single-authored papers. Other studies have assessed women authorship, and found that a gender gap in published literature persists. This study applies the large sample size and methodology of West et al. 2013 to the broad discipline of aquaculture, and compares these results to gender authorship in the International Aquaculture Curated Database (IACD) – a compilation of 543 peer-reviewed publications supported by four international aquaculture programs headquartered at Oregon State University -- and two curated databases in the JSTOR in the Web of Science.

Results reveal that the percentage of women authors (13.8%) was similar for the JSTOR aquaculture subsample and the IACD (15.7%), yet significantly lower for that of the Web of Science database (3.7%). Women are not well represented any of the databases, and remain underrepresented as authors in any position in aquaculture journals. To contextualize our findings, we examined the number of women graduates in agricultural, biological, natural, and social sciences who earned degrees in the U.S. from 1991-2015. Results from the U.S. Department of Education’s National Center for Education Statistics and the percent of female graduates in the IACD show that the percent of women graduates each year has increased with women representing more than 50% of graduates, providing contextualization for the proportion of women in the discipline. Learning how authorship has changed in the aquaculture discipline over the recent decades is critical for promoting gender equity for future aquaculture scholarship and the sustainability of the professional discipline.

## INTRODUCTION

Studies have found that women are underrepresented in science, publish less (Martin 2012; Conti and Visentin 2015), and receive less grant funding than their male counterparts (Vernos 2013). Other studies have assessed women’s authorship in disciplines including political science and medicine, and found that not only does a gender gap in published literature still remain, but women’s authorship has been levelling off in recent years (Breuning and Sanders 2007; Jagsi et al. 2006; and Dubey et al. 2016). From examining authorship of more than eight million papers across disciplines in natural sciences, social sciences, and humanities, West et al. (2013) found that men dominate in the first and last authorship positions and that women are underrepresented as single authors. These numbers matter because authorship position, first and last typically getting the most credit, is a major component of university evaluations of researcher proficiency. This criterion is applied to determine promotions, assessments for tenure-track positions, attainment of research funding, and so on. Therefore, authorship position can be used as a proxy for the status of gender integration and diversity in academia.

The problem with relying too heavily on authorship position for evaluating a researcher’s success is that there is no straightforward process across disciplines for assigning authorship order. The process of determining each author’s contribution to a paper and assigning authorship position varies across academic institutions, disciplines, and sub-cultures within research groups. This is partly because it can be difficult to ascertain how much work each contributor has put into a paper (Laurance 2006; Tscharntke et al. 2007). Traditionally, the first author has contributed the most to the paper and receives the most credit, and the positions of the subsequent authors are determined according to contribution, alphabetical order, or reverse seniority (Tscharntke et al. 2007). The last author often gets as much credit as the first author, as they are assumed to be the intellectual or financial driving force (Tscharntke et al. 2007).

Subtle biases and other factors can influence how authorship is assigned. Increasingly, “gift authorships” are given, i.e., an author is added for courtesy reasons because of their academic status, particularly in biomedical journals. This trend further confuses the actual contribution of each author listed on a publication. Because of the unclear process by which the set of authors for a paper is determined, identifying the amount of work each author contributed is challenging. The culture of peer-reviewed publications is also changing and this also affects how changes in gender authorship over time are assessed. In particular, over the last several decades, the amount of collaborative and cross-disciplinary research has grown, as has the pressure to publish. Both of these factors have led to growth in the number of authors listed per paper (Wren et al. 2007). The growing number of authors per paper makes it even more difficult to adequately and fairly assert authorship order.

While studies have revealed gender inequities in authorship in scholarly literature, no such study has been completed for the aquaculture discipline. The academic discipline of aquaculture is relatively new and interdisciplinary, and many aquaculture degrees are granted from fisheries departments. Our analysis of the discipline, therefore, is embedded within the broader domain of fisheries. In more than 50 academic institutions, a study by Arismendi and Penaluna (2016) found that women and minorities are still a small portion of tenure-track faculty in the discipline of fisheries. Over the past three decades, they found only a slight increase in the inclusion of women among the academic community of fisheries science. This suggests a perpetuation of the “leaky pipeline” in fisheries science as, in recent years, women have received more than half of the doctoral degrees in the biological sciences (Miller and Wai 2015; Egna et al. 2012; Blickenstaff 2005). These trends and a study by Penaluna (2005) reveal that women are less likely to be promoted than men in academia, and the unlikelihood of a promotion can be linked to the status of gender authorship in peer reviewed literature. Ignoring these inequities or allowing them to persist limits the development of the scholarly field of aquaculture. By attempting to conduct a gender authorship analysis for aquaculture, we’re helping to promote the development of the fastest growing food production sector in a relatively new and interdisciplinary scholarly discipline. A better understanding of gender integration in the discipline is the first step in understanding how to overcome barriers to the sector’s growth.

This study evaluates the status of gender authorship in aquaculture by comparing authorships across the JSTOR Corpus database archive to a subsample of JSTOR and the Web of Science with aquaculture journals, and to a smaller, curated database, compiled by the Aquafish Innovation lab, of aquaculture peer-reviewed publications. The International Aquaculture Curated Database (IACD), was created in order to have a very rich data source of aquaculture publications from around the world that have been published throughout the entirety of the existence of modern era of aquaculture for scholarly analysis. The richness of an international curated database lends itself to factoring in additional variables such as funding and faculty rank, along with other social metrics when assessing authorship. The present paper shares findings that the percentage of women authors across the aquaculture discipline is significantly lower than women’s apparent presence in the discipline. Since women have received more than half of the doctoral degrees in the biological sciences, it is plausible that women represent more than 16% of researchers working in the discipline, while this is the rate at which women are authoring papers. This number is corroborated across two completely disparate, yet valuable sources within the discipline.

## MATERIALS AND METHODS

In order to build on the work of West et al. (2013) and other similar studies conducted on gender authorship in peer-reviewed literature for the aquaculture discipline, we compared multiple data sets. The first and richest dataset, the International Aquaculture Curated Database (IACD), was built by the AquaFish Innovation Lab, and consists of 543 articles written by 1706 authors in 121 journals, all of which were published between 1983-2016. The IACD draws from peer-reviewed papers whose research was supported by four separate international aquaculture programs, which were developed by Hillary Egna including: (1) Pond Dynamics/Aquaculture Collaborative Research Special Program (CRSP) (1982-1996); (2) Aquaculture CRSP (1996-2008); (3) AquaFish CRSP (2006-2013); and (4) AquaFish Innovation Lab (2013-Present). AquaFish Innovation Lab staff reviewed both electronic and hard copies of journal articles, including full names, gender of authors, and author position, with the percentage of unknowns being less than 1%. The gender of the authors was recorded by Egna from having a personal connection to the author or by the lead authors themselves.

The IACD was analyzed for comparison to three other datasets: two separate JSTOR collections (The Recalibrated JSTOR and the JSTOR Subsample) and a Web of Science dataset. JSTOR is an expansive database of publications organized according to broad topics, and contains publications dating back to 1665, and was used for the West et al. (2013) authorship study. Web of Science is a similar online database, as well as Academic Search Premier, Scopus, and Microsoft Academic Graph (MAG). Each database has proprietary strengths and weaknesses. JSTOR has far more time depth than any of the other databases and it has full text for all their articles whereas most of the others have only bibliographic data. Web of Science has decades of data. Hundreds of databases have been created, but many of them are specific to certain disciplines or types of publications, whereas those listed above are more comprehensive across the literature. By comparing the IACD to both JSTOR and the Web of Science, more journals within the interdisciplinary discipline of aquaculture are captured in this analysis. JSTOR Re-calibration was done in order to revisit the gender findings from West et al. (2013) and compare the findings to authorship data in the present study. The JSTOR aquaculture subsample separated the aquaculture journals from others within the broad database. It begins in 1913 as that was the year one of the first aquaculture-related journals began. JSTOR journal areas include: cultural studies, arts, business and economics, history, humanities, law, medicine and health, science and mathematics, and the social sciences. Aquaculture journals are located within the science and mathematics category. Web of Science is an online subscription-based scientific citation indexing service produced by the Institute for Scientific Information and includes science, social science, arts, and humanities disciplines. From the more than 90 million records, we extracted articles from more than 100 journals within the aquaculture discipline. This includes all of the journals in the IACD plus more that are commonly publish research in aquaculture.

In the JSTOR and Web of Science, authorships are defined as an author-paper relationship, and does not count unique authors. This requires author disambiguation for the full databases, which is an ongoing challenge in the field of bibliometrics and scientometrics. Because of the large number of authorships in JSTOR and Web of Science, gender was inferred by looking up the frequency of first names in the U.S. Social Security Database. For example, if “James” appears 99% of the time as a boy, we assume that an author with the name “James” is male. For androgynous names such as “Andrea” and first names written as initials, we could not infer gender so we do not include these authors in the analysis. Therefore, the gender labels are self-identified and determined by only looking at the names and the frequency of gender for a given name. Unidentifiable names account for about 1 in every 5 authors in the Recalibrated JSTOR dataset (Table 1).

**Table 1.** Four datasets used for this study with varying journals, articles, authorships, time periods, and percent of genders unknown.

| Dataset | #Journals | #Articles | #Authorship | Time<br>Period | % Genders<br>Unknown |
| --- | --- | --- | --- | --- | --- |
| JSTOR | 2227 | 1.8 million | 2.8 million | 1666-2011 | 26.7% |
| JSTOR Sub-<br>sample | 8 | 23,381 | 43,146 | 1913-2016 | 23.7% |
| IACD | 121 | 543 | 1706 | 1983-2016 | <1% |
| Web of<br>Science | 185 | 494,531 | 496,745 | 1980-2016 | 69% |

The Recalibrated JSTOR Corpus and Web of Science cover all major realms of scientific publications; the aquaculture subsample of the JSTOR Corpus and the Web of Science include a large number of articles from a select few aquaculture journals; and the IACD is a substantiated aquaculture-specific database containing fewer journal articles. The IACD, JSTOR, and Web of Science comprise journals in the biotechnical domain of aquaculture more so than in the social or management domains of the discipline. Together, the four data sources allow for a stronger understanding of gender representation in journal authorship.

Lastly, we contextualized the findings from these datasets within the percentage of women graduating in aquaculture, as well as factored in how the field has grown over time. As aquaculture degrees were not conferred widely or until recently in academia, assumptions were made to cover the wide range of academic disciplines that could relate to aquaculture.

**Figure 1.**
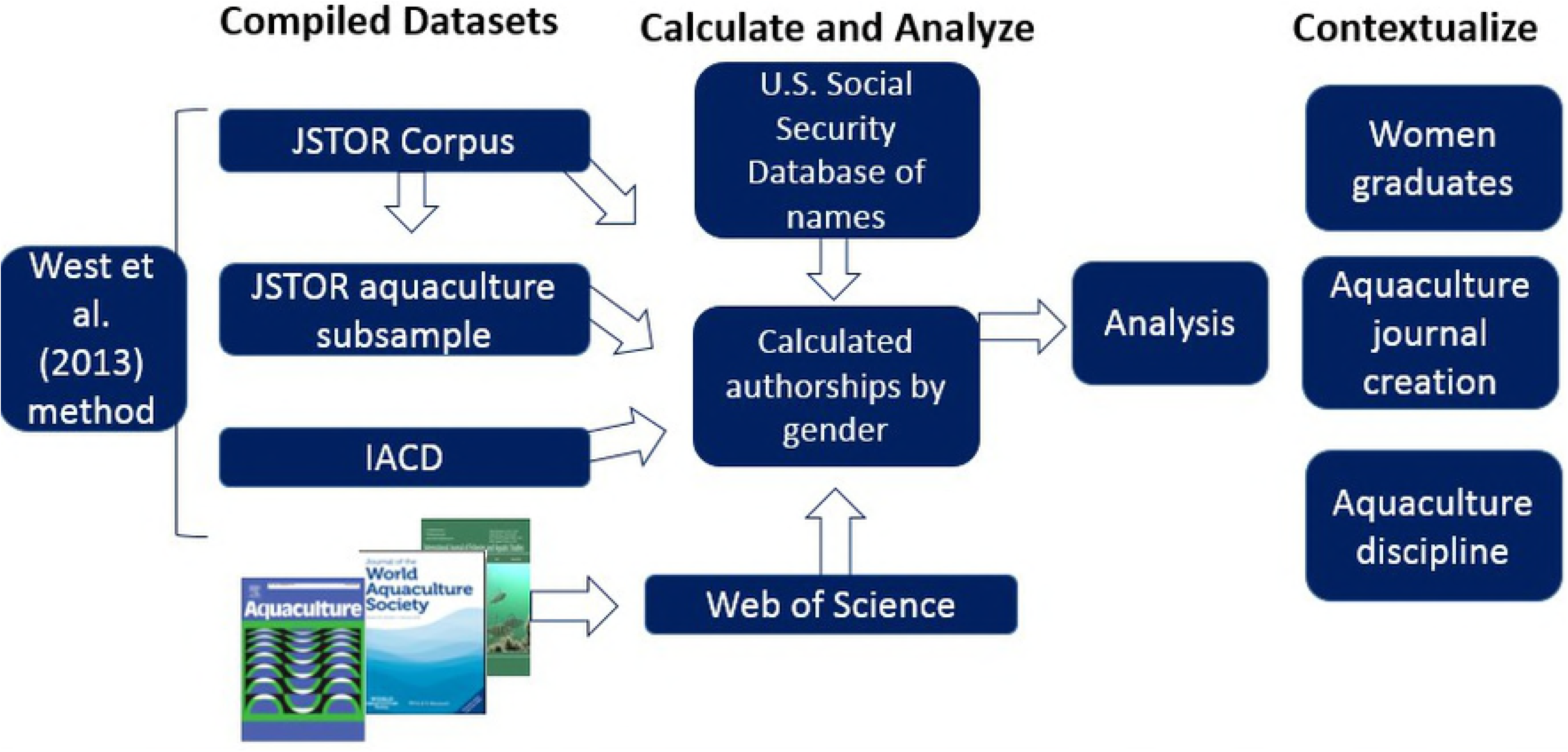
Flow chart of methodology used for this study.

## RESULTS

In the entire JSTOR Corpus, comprising nearly 2 million papers, women represent 21.9% of total authorships for papers published between 1665-2011 (West et al. 2013). This timeframe represents the existence of JSTOR publications. For comparison, in fisheries-related fields such as Ichthyology and Aquatic Ecology, women represent 21.0% and 9.0% of total authors, respectively. In the JSTOR aquaculture subsample, 23,381 articles and 43,146 authorships within eight aquaculture journals (since 1913) were extracted and assessed for authorship gender in multiple positions to compare to the Recalibrated JSTOR dataset. The JSTOR recalibration adjusted for the period in which the first aquaculture journal in our subsample was initiated. The following eight journals were selected because they are highly ranked in the aquaculture discipline: Ambio, Copeia, Estuaries and Coasts, Journal of Coastal Conservation, Journal of the North American Benthological Society, Limnology and Oceanography, and Water and Environment Research. We recognize that these journals do not comprise a representative sample of all aquaculture journals, and are skewed towards biotechnical domains of aquaculture. However, these journals are consistent with the journals available in JSTOR. In the Web of Science, comprising almost 500,000 articles in the subsample extracted for this study, women represent 8.5% of the total authorships for papers published between 1980-2016. This timeframe is in line with the IACD for comparison. This analysis includes articles from 185 journals that are considered relevant to the aquaculture discipline.

Table 2 outlines our findings by significant authorship position across the four main datasets of peer-reviewed aquaculture literature. Across the board, women represent between 9-15% of significant authorship positions in these four datasets. Due to the methodology of assigning genders to author names within the JSTOR alongside the U.S. Social Security Database, there are higher percentages of unknown genders for the JSTOR and Web of Science datasets than for the IACD. Results show that women occur in low percentages as authors in any position in aquaculture journals, reinforcing results found by West et al. (2013) more generally in science. Women represent 16.1% of authorship in all positions in the Recalibrated JSTOR Corpus and only 8.5% in the Web of Science, after correcting for unknowns. The percentage of women authors was comparable for the JSTOR aquaculture subsample (13.8%) and the journals in the IACD (15.7%), but much less so for the Web of Science (Table 2). For single-authored papers, the JSTOR Corpus shows an overall decline over time. However, there has been an increase in sole authorship by women. Before 1990, only 12% of single authored papers were written by women. After 1990, that number increased to 26%. In the JSTOR aquaculture subsample, women represent 11.0% of single-authored papers since 1913, and 17.7% in the Web of Science, respectively. In the IACD, women represent 11.1% of all single authored papers since 1990.

**Table 2.**
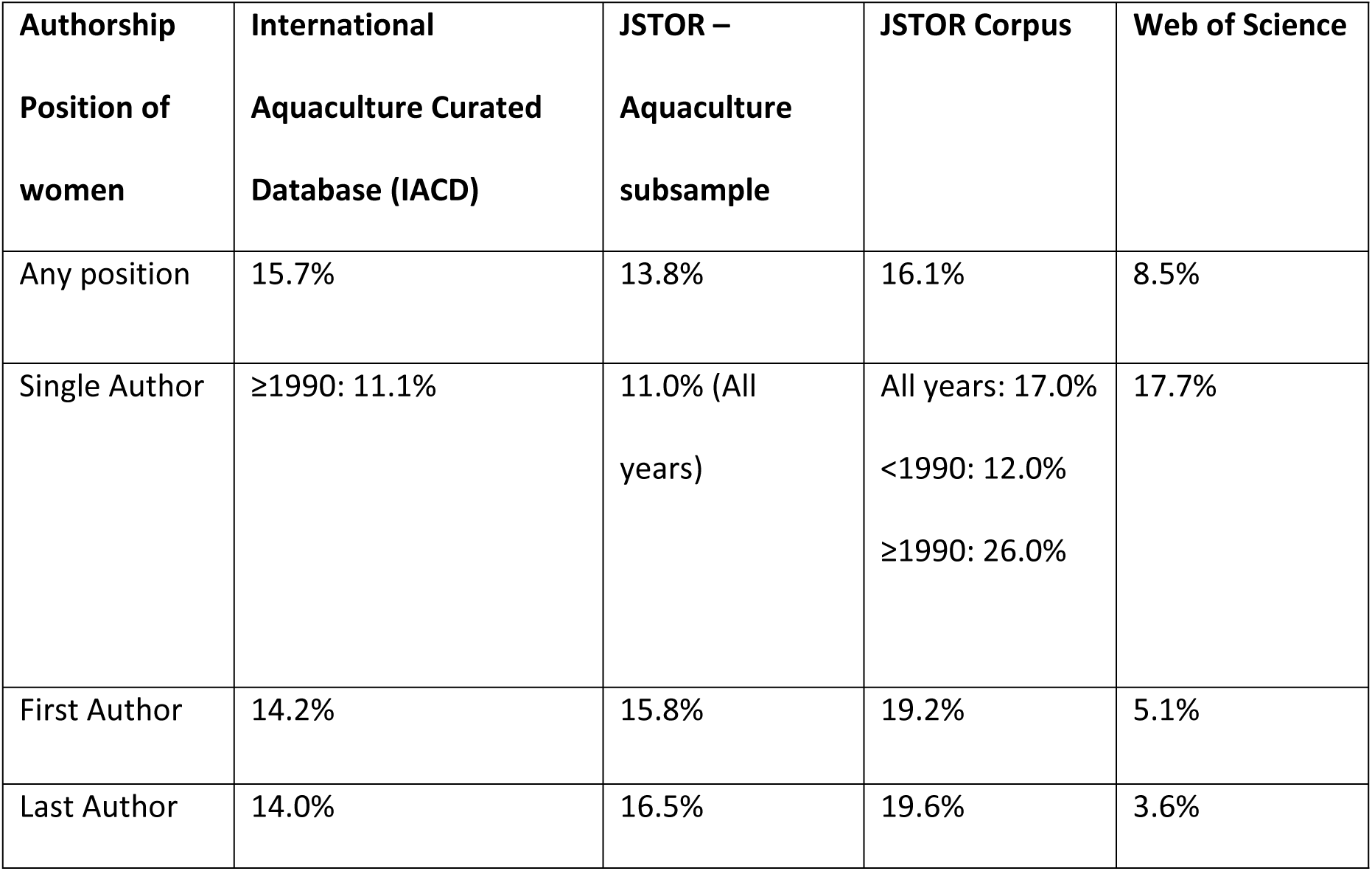
Findings by significant authorship position across the four datasets.

Percentages of women in last authorship positions were comparable for the publications in the JSTOR Aquaculture subsample and the IACD at 15.8% and 14.4%, but were much less for the Web of Science at 3.6%. A similar trend is seen with first authorship positions where the JSTOR Aquaculture subsample are comparable at 15.8% and 14.2%, while the Web of Science is only a mere 5.1%. First and last author results from the overall JSTOR Corpus for all fields were slightly higher than for the field of aquaculture at 19.2% for first authorship and 19.6% for last.

As well as recent changes in the publication process for peer-reviewed literature, the history of aquaculture was considered for this analysis. To understand the evolution of gender in the aquaculture discipline, it is important to first recognize that the discipline of aquaculture has changed substantially over the past 30 years (FAO 2016). Global aquaculture production took off in the early 1980s, and rapidly expanded through the 1990s to present to accommodate a growing global population with its changing diets and preferences. Development was especially expansive in the 1980s, with pond culture predominating total aquaculture production. The fisheries discipline has also grown in both scope and geographic range. There has been a global scale expansion of marine fisheries from the North Atlantic and West Pacific to the Southern Hemisphere. The southward expansion of intense industrial fisheries exploitation occurred at a rate of almost one degree latitude per year with the greatest expansion occurring in the mid-1980s and early 1990s (Swartz et al 2010).

Growth of the aquaculture discipline and industry have, not surprisingly, mirrored each other. Preliminary data from over 300 aquaculture-related publications shows the rapid inception of new journals from the late 1980s to the 2000s. Overall, the number of journals and publications has grown in all disciplines. In the Recalibrated JSTOR set, we find that roughly half of all peer-reviewed publications were published after 1990. We think that this is consistent across other large scholarly article corpora. Scientific publishing, like many other industries, has faced many changes with the onset of the internet. Journal articles today are accessed online with increasing frequency, and retrieved in digital formats rather than through printed sources (Laakso et al. 2011). The way that journal articles are accessed online has also changed in recent years, particularly with the growth of Open Access publishing between 1993-2009. Since 2000, the annual growth rate for Open Access journals has been 18%, and 30% for the total number of published articles (Laakso et al 2011). The evolving mechanisms for publishing peer-reviewed literature have consequences for researchers in the field, and their authorship track records.

Figure 2 shows the years that major aquaculture journals began (n=166). There was significant growth in aquaculture journals in the early 1970s through the 1990s. For example, JWAS began in 1970. While this is not a comprehensive list of all of the journals that ever publish aquaculture articles, it represents most of the major journals in the discipline that had initiation years available online. Figure 2 follows a similar curve to that of the global aquaculture production, which started to increase in the early 1980s, and began rapidly expanding in the 1990s to the present to accommodate a growing global population. The discipline has growth both in scope as well as geographic range.

**Figure 2.**
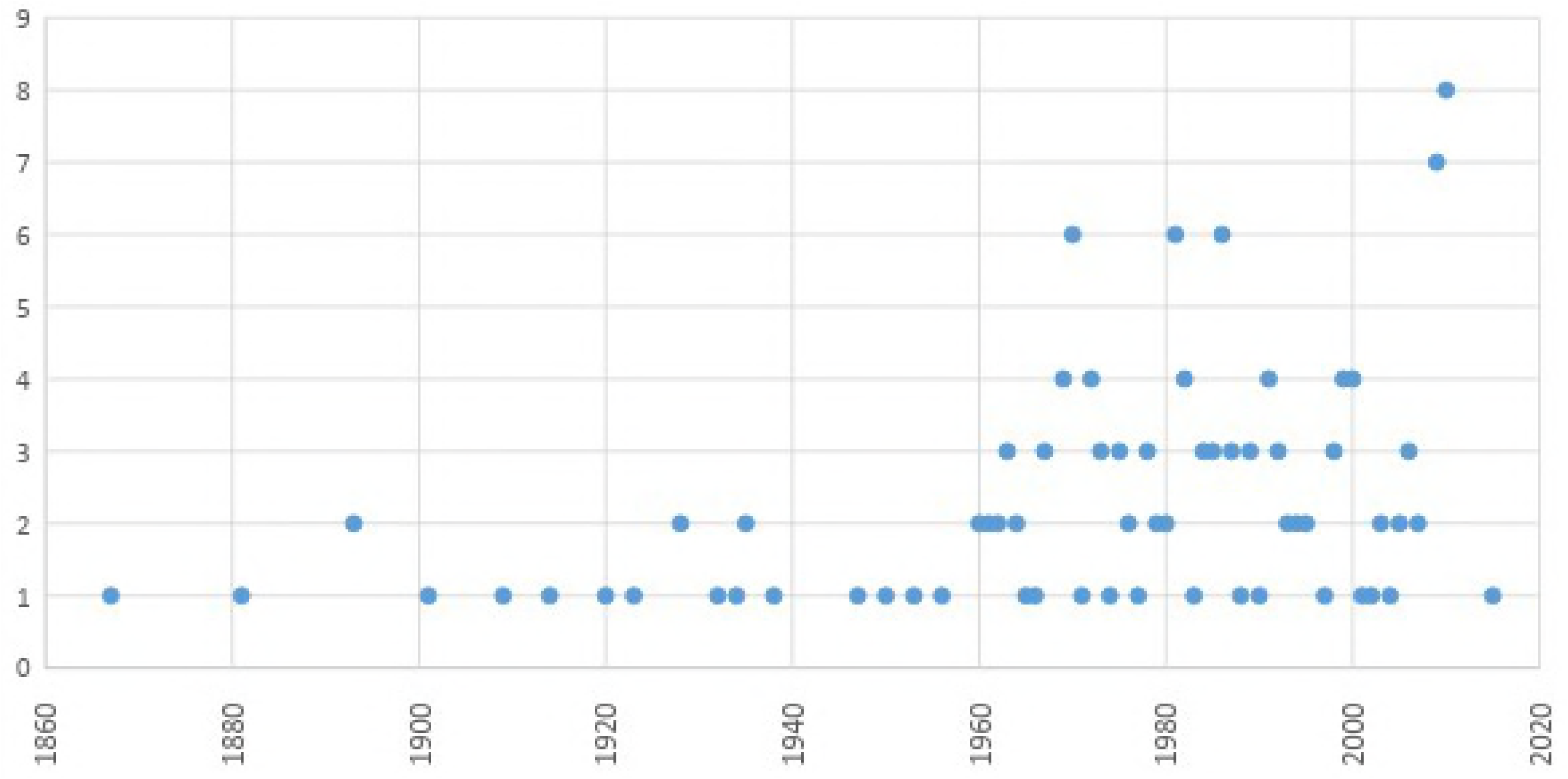
History of aquaculture and journal initiation over time.

Figure 3 shows the percent of each position in the IACD for each year between 1990-2016. Men first and last authorships dominate the journal articles published each year, with women single authors being the lowest. However, the gap between men and women authors does seem to decrease over time, which leads us to believe that women’s status in the field is improving.

**Figure 3.**
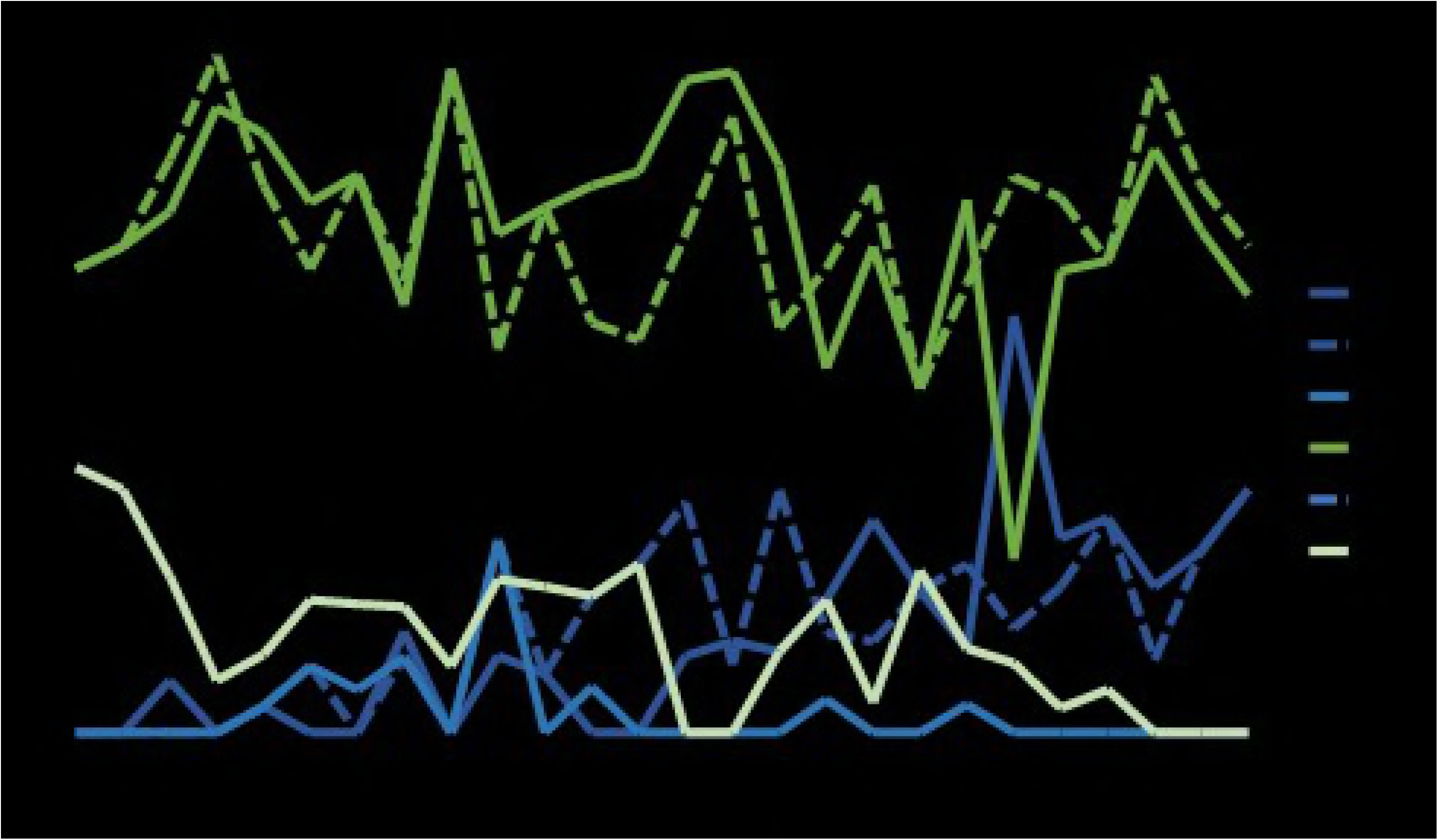
Women authorship by position over time in IACD.

To contextualize our findings with the percentage of women graduating in the field, we examined several sources to better understand the numbers of women graduates in aquaculture. Because of the relatively nascent, and interdisciplinary nature of aquaculture, we applied sources from within the U.S. and international as well as across disciplines including fisheries, biological, agricultural, and social sciences. According to Elsevier, approximately 28% of researchers around the globe are women, yet only 13% of highly cited authors in 2014 were women (Elsevier 2015). In the U.S., we used the U.S. Department of Education’s National Center for Education Statistics to quantify the number of female graduates in agricultural, biological, natural, and social sciences who earned Bachelor’s Master’s and PhD’s in the U.S. from 1991-2015. Figure 4 shows the percent of female graduates each year at each degree level. The proportion of women graduating in the field has increased over time, with the most obvious increase being that of PhD graduates, representing roughly 30% of graduates in 1991 to more than 50% of graduates in 2015. Additionally, Arismendi and Penaluna’s 2016 study on women publishing in fisheries, found that women and minorities are still a small portion of tenure-track faculty in the discipline of fisheries. Lastly, we evaluated the percent of women AquaFish graduates per year, and found a slight increase over time, with no significant upward trend.

**Figure 4.**
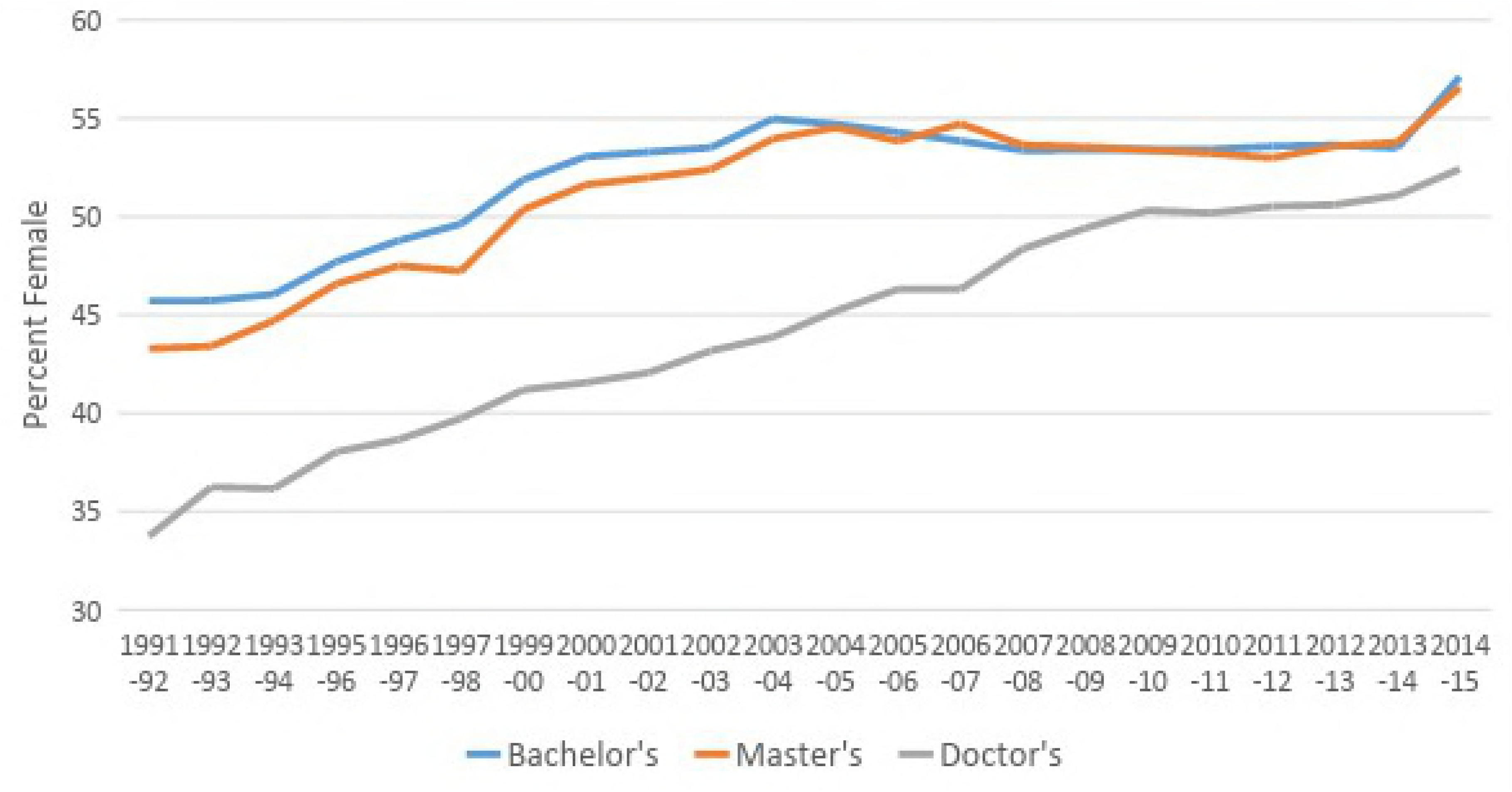
Percent women graduates in Agricultural, Biological, Natural and Social Science. Source: U.S. Department of Education, National Center for Education Statistics.

This analysis is very useful as many students publish their research chapters soon after they graduate, despite whether or not they continue to work in academia and publish. While these graduates do not represent all of the women science graduates internationally, since the data is U.S.-based, it is a still a useful comparison for a general understanding of how many women are graduating in the agricultural, biological, natural, and social sciences, all of which feed into aquaculture scholarly literature.

Since the IACD percentages reflect that of the JSTOR sub-sample and Corpus, it is a proxy for the women authors in the discipline as compared to women graduates in science in the U.S. Figure 4 shows percent women graduates with Master’s and Phd’s in agricultural, biological, natural, and social sciences from 1991-2015, in black, alongside the percent of significant authorship positions women held each year for the IACD. These numbers are from the U.S. Department of Education’s National Center for Education Statistics. There is a slight increase in women authorship positions as a percentage of the total publications for each year, over time.

**Figure 5.**
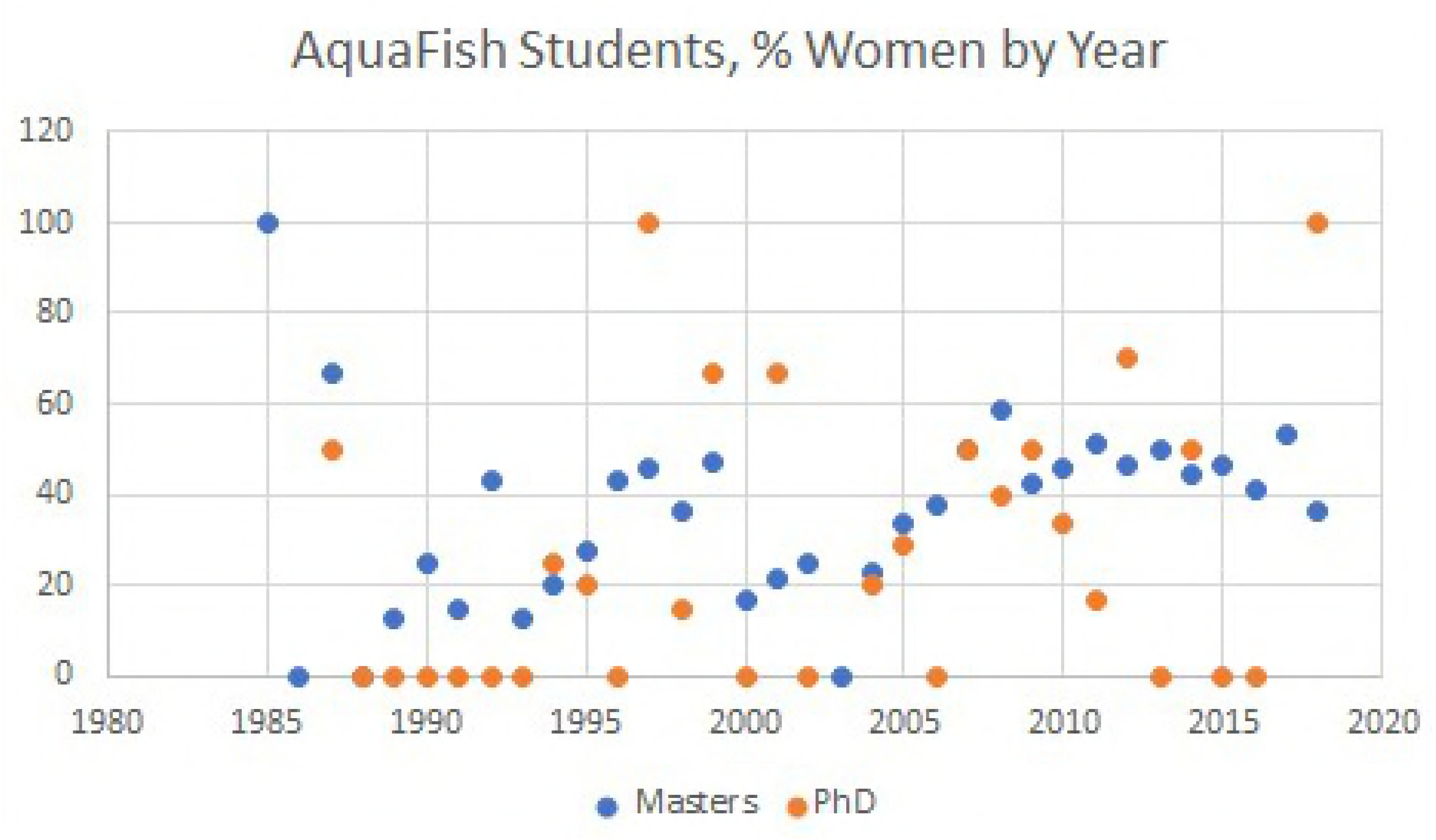
AquaFish graduates as percent women by year.

**Figure 6.**
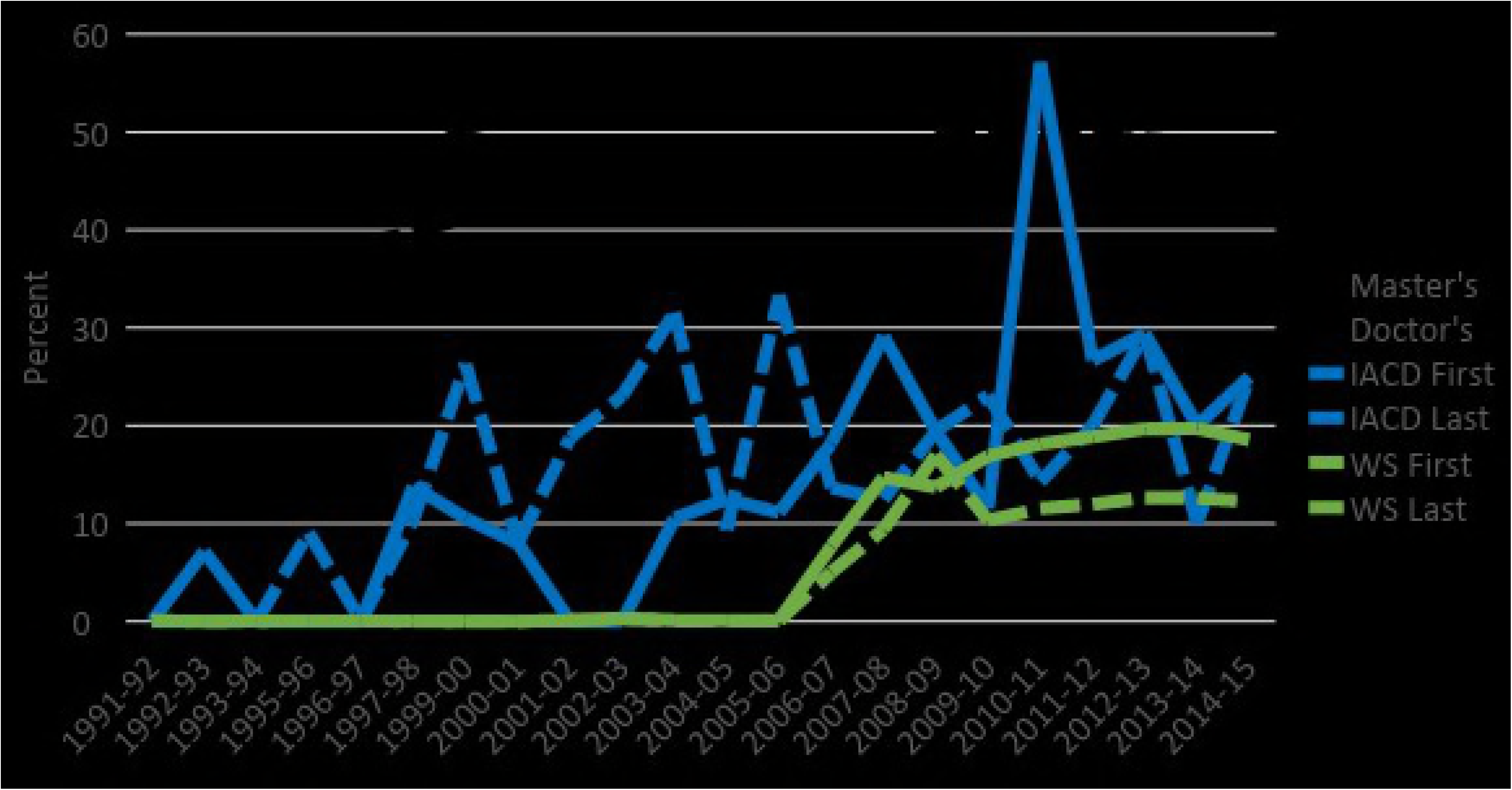
Percent women graduates in science alongside percent first and last authorship positions in the IACD and Web of Science datasets.

## DISCUSSION

It appears that the gap in women authorship is closing, however women authorship remains low considering the increasing proportion of women graduates in aquaculture sciences. However, the U.S. data does not represent the proportion of women that are actively publishing in aquaculture as an academic discipline. Moving forward, it is important to encourage organizations and individuals to consider how structures that propagate gender bias can be overturned to promote better outcomes in authorship, hiring, and promotions.

These findings can be applied to the greater context of women in academia. In 2015, Elsevier published a study of research performance through a gender lens across 20 years, 12 geographies, and 27 subject areas to share insights and guidance on gender research and equity policy with governments, funders, and institutions worldwide. They found that approximately 28% of researchers around the globe are women, with only 13% of highly cited authors in 2014 were women. However, there is a drop off in degrees, starting at the PhD level. Further, health and life science have the highest representation of women among researchers. Studies like Elsevier’s are continuing to explore why the leaky pipeline occurs, and why women are dropping out of academia in their PhD.

Given this study and others, we recommend a number of steps to combat gender inequity in aquaculture scholarly literature and other academic disciplines. First, it is important to continue to track authorship to measures success or weaknesses in progress towards integration. Standardized practices for assigning authorship position would be mainstreamed and made transparent. Faculty and mentors should encourage women scientists to remain in academia through mentoring, opportunities for promotion, and opportunities to review and edit other publications. While we do not yet have details on non-unique identifiers for prolific authors and people with multiple degrees, this could be an important next step to better understanding trends in authorship position by gender.

## CONCLUSION

Comparing the percentage of women authors across all four databases reveals a low percentage of women authors -- between 8.5% --16.1% of all authorships. The four data sets represent a wide range of aquaculture journals that are well regarded within the discipline. These results for aquaculture echo the findings of West et al. (2013) for women in many fields of science, as well as (Arismendi and Penaluna 2016) on the status of women publishing in the broader discipline of fisheries.

While there are many factors that may explain why women hold a low percentage of authorships across all fields of peer-reviewed literature and in aquaculture, in particular, these results do not reveal the cause. The data reflect an end-result that is influenced by a number of factors that are not easily studied and have not yet been addressed in the project. One of the main factors is the proportion of women trained and actively working in the aquaculture discipline. Also, recognizing that gender is a social construction, our preliminary work was simplified by binary designations (man-woman; male-female); additional deeper analyses may reveal nuances for other underrepresented groups.

Since it is known that women have been reported by the World Bank (2008) to comprise 47% of the total workforce in fisheries, this is a rough estimate confounded by a paucity of gender-disaggregated data in aquaculture and fisheries overall. Few data are available on the percentage of women in the fisheries discipline. One exception is the study by Arismendi and Penaluna (2016) for the United States of America. In that study, 26% if federal fisheries scientists and managers, and 31% of research faculty were women. Until adequate numbers for women in aquaculture and in the aquaculture discipline are obtained, it is useful to apply information from the greater field of fisheries to frame the research.

These results suggest that gender inequities in aquaculture, specifically in authorship of peer-reviewed literature, exist. While these are general conclusions, 15% is a relatively low number for women authorships in aquaculture considering that the proportion of women authorships in the entire JSTOR corpus is 22%. The IACD may prove a useful tool for social network analyses including assessments of unique very highly networked authors, and of subsequent generations of authorships. The richness of an international curated database lends itself to factoring in variables such as funding and faculty rank, along with other social metrics. The information in these data sets can be used by other studies to assess the major influences on gender equity in the field of aquaculture. Increasing awareness of the equitable treatment of scientists in aquaculture remains essential for the sustainable growth of the discipline.

## Acknowledgements

All listed authors contributed a significant amount to the paper. Dr. Hillary Egna had the original intellectual contribution to the work and a strong vision for the paper. She also contributed to data collection, analysis, writing and editing. Morgan Chow collected the IACD information, analyzed results with the JSTOR databases, and wrote the backbone of the paper. Dr. Jevin West conducted data analysis for the recalibrated and subsample of JSTOR, while providing substantial information for the methods. AquaFish Innovation Lab intern, Katie Nye Hogen assisted with IACD data collection and compilation. This research is a component of the AquaFish Innovation Lab, which is supported in part by the US Agency for International Development (USAID CA/LWA No. EPP-A-00-06-0012-00), and in part by participating institutions. The AquaFish accession number is 1464. The opinions expressed herein are those of the authors and do not necessarily reflect the views of the AquaFish Innovation Lab or USAID.

